# Blocking CCR2^+^ Monocyte Infiltration Enhances Neurological Recovery after Subarachnoid Hemorrhage

**DOI:** 10.1101/2025.03.18.644055

**Authors:** Dong su Kang, Seung Hyeok Seok, Seung-Hoon Lee, Yirang Na

## Abstract

**BACKGROUND:** Subarachnoid hemorrhage (SAH) patient’s mortality rate has been decreasing, but improving survivor’s prognosis outcome, which impacts patient’s quality of life, is still limited. An Early Brain Injury (EBI), which is a period of first 72 hours after SAH, is critical phase when resident and infiltrating innate immune cells induce neuroinflammation, deciding the prognosis outcome. While the roles of microglia have been studied, little is known about the role of infiltrating immune cells, especially that of monocytes. We investigated how the absence of monocyte infiltration during EBI may affect SAH outcomes.

**METHODS:** Using Middle cerebral artery (MCA) perforation method, we established Wild type (WT) and C-C Chemokine Receptor type 2 (CCR2) knockout transgenic mice (CCR2^-/-^) SAH model. Using flow cytometry, we determined differences in immune cell infiltration population. Also, through Cytometric bead array (CBA), we measured inflammatory cytokine level. We performed different behavioral experiments to assess neurological and behavioral recovery and prognosis of SAH. Furthermore, to confirm neuronal cell death severity, immunohistochemistry was performed. We also tested CCR2 antagonists in the early EBI period to see if it impacted WT mice’s neurological outcome.

**RESULTS:** SAH model showed increased infiltration of CD45^hi^ immune cells, mainly Ly6C^+^ monocyte at 24 hours after SAH, which were nearly abolished in CCR2^-/-^ SAH mice. Increased IL-6 and TNF-α cytokine levels in cerebrospinal fluid was significantly reduced as well. Also, better neurological and motor recovery was observed from CCR2^-/-^ SAH mice. Overall neuronal cell death by late EBI period was significantly decreased. Finally, WT SAH mice treated with CCR2 antagonists showed improved neurological outcomes 24 hours after SAH, while showing reduced total infiltrating immune cells and monocytes.

**CONCLUSION:** Classical monocyte infiltration, which occurs via CCR2 signaling, is detrimental for neuroinflammation during EBI of SAH that leads to poor prognosis. CCR2 inhibition could be a potential target for interventional therapeutic strategy.

Graphical Abstract

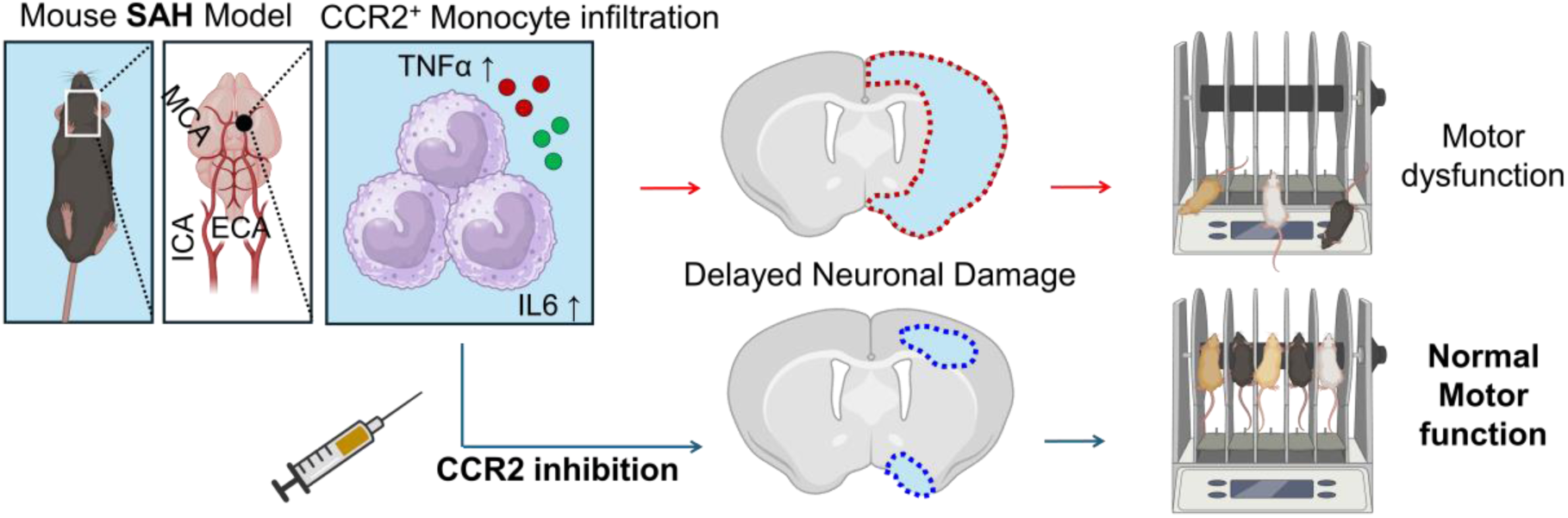

## Introduction

Subarachnoid hemorrhage (SAH), a hemorrhagic stroke that occurs due to traumatic brain injury or vascular abnormalities such as aneurysms, is a critical pathology for its high fatality rates and secondary complications^1–3^. Even though mortality rates have been decreased due to advanced surgical interventions^4^, pathophysiological burdens of surviving patients such as vasospasm, delayed cerebral ischemia (DCI), and long-term neurological deficits remains a burden^3,5,6^. While vascular complications have available countermeasures such as nimodipine^7,8^, effective alleviations of neurological complications like motor weaknesses are limited.

Recent studies suggest Early Brain Injury (EBI) of SAH, which refers to 72 hours after the initial SAH, is a critical time that determines severity of such neurological outcomes^9–11^. EBI comes with several pathophysiologic features such as increased intracranial pressure (ICP), vascular endothelial damage, and microvascular dysfunction^12–14^. One key features of EBI are the neuroinflammation, which induce high fever and neuronal cell death^15–17^. While neuroinflammation during EBI and their importance in prognosis has been recognized, roles of different immune cells involved in this neuroinflammation are understudied. Neuroinflammation in SAH commonly occurs in response to extravasated blood^17^. Typically blood in subarachnoid space initiate microglial activation^18–20^, and infiltrating myeloid cells are recruited, elevating the neuroinflammatory response during EBI^21^. One of the main immune cells recruited at the site of injury are Ly6C^+^CD11b^+^ classical monocytes, which moves through C-C Chemokine Ligand 2 (CCL2 or MCP-1) and receptor (CCR2) signaling^22^. While the presence of these infiltrating monocytes is definitive, how their presence impacts neuroinflammatory response and neurological outcome remains unclear.

In this study, we used CCR2^-/-^ transgenic mice to prevent monocyte infiltration, and induced SAH model to investigate the role of infiltrating classical monocytes. We report prevention of CCR2^+^ dependent monocyte infiltration results in decreased cytokine levels during EBI period of SAH, along with decreased overall immune cell infiltration during the first 24 hours of SAH. We also show significantly improved neurological recovery CCR2^-/-^ SAH mice model compared to WT, with conserved motor function. We also found reduced neuronal death as well. Finally, CCR2 inhibition drug treatment in early EBI period results in amelioration of WT mice’s neurological scores. Overall, our evidence suggests that CCR2^+^ monocytes play an important role during EBI period of SAH and may be an important therapeutic target for amelioration of its prognosis outcome.

## Methods

The authors declare that the original data in this study are available upon reasonable request.

### Animals

All animal experimentations and animal care were carried out following the guidelines approved by the Institutional Animal Care and Use Committee (IACUC) of Seoul National University (IACUC No. SNU-240220-1). Mice were housed in the animal care facility, under 12-hour light/dark cycle with access to food and water. All possible measures were taken to minimize suffering and reduce the number of animals used in this experiment. All mice used for modeling were within the age of 12 to 14 weeks, which allowed easier access to Internal Carotid artery (ICA) for MCA perforation modeling. All mice had a C57BL/6 background, and the following strain was used as WT or control. C57BL/6-Tg (CD68-EGFP)1Drg/J was used for meningeal wholemount imaging. B6.129S4-Ccr2^tm1Ifc^/J Homozygotes were selected for the use of all CCR2^-/-^ SAH models.

### SAH modeling

To create SAH model, we followed the SAH modeling protocol from Buhler et al.,2015^23^ with slight modifications. All mice prior to surgery were under anesthesia using 4 % isoflurane in 30 % oxygen and were maintained using the anesthesia systems (Vetflo). Mice were laid in supine position with exposed neck, and skin was opened near slightly off the mid-line towards the ipsilateral side of targeted side. Connective tissues were bluntly dissected through until the external, internal, and common carotid artery (ECA, ICA, and CCA respectively) were identified. Using a black silk 6-0 filament, ECA is ligated and CCA was temporarily ligated with silk line and hemostats. Stiffened, sharply cut Blue Nylon (5-0) was used for perforation, which occurred initially from ECA, traveling through ICA until it reached the MCA. After the MCA perforation, all mice after SAH modeling was placed in a cage with Infrared heat for 20 minutes, or until they awakened from anesthesia to prevent hypothermia.

### Meningeal Wholemount

For Meningeal wholemount to confirm immune cell infiltration, we followed the protocol in accordance with Louveau et al.,2019^24^ with slight modifications. First, Skull cap of C57BL/6-Tg (CD68-EGFP)1Drg/J (hCD68-GFP) mice were harvested after euthanization. Then, Skull caps were fixed in 4% PFA in PBS overnight at 4°C. Then under a petri dish and a dissecting microscope, meninges were carefully detached. Detached meninges were washed with PBS for 5 minutes and went through the same protocol written under Immunofluorescence staining.

### Flow cytometry (FCM)

For determining immune cell population, Flow cytometry was performed. Mice were anesthetized and perfused with PBS prior to harvesting the brain. Prior to the brain extraction, Cerebro Spinal Fluid was obtained from their Cisterna Magna using an insulin syringe (BD). Brainstems and cerebellum were removed prior to harvesting. Samples were minced <1mm in a collagenase V (Sigma, C9263, 1mg/ml) infused with 5% FBS in PBS with DNase (100unit), incubated for 20 minutes for single cell dissociation. Next, a sample was given EDTA to stop the reaction of digestive enzyme and went through 40μm cell strainer into 50 ml tube. Cells were collected by centrifuge of 1300rpm x 5 min. Immune cells were collected using 30% Percoll™ density gradient media (Cytiva) in 1xPBS and removed all myelin. Then, 6ml of Ammonium-Chloride-Potassium (ACK) Lysing buffer (4M Potassium bicarbonate, 0.5M Ammonium Chloride in Triple Distilled Water) was applied for 5 minutes to remove any red blood cell remaining. The ACK buffer was stopped by adding 1xPBS up to 50ml, then were centrifuged at 1300rpm for 5 minutes. Antibodies of the following were diluted 1:500 ratio in FCM buffer (5% FBS in PBS, filtered, with 0.1% Sodium Azide) and stained for 20 minutes on ice: CD45 BUV395, CD11b V450, Ly6G PE, CX3CR1 APC Cy7, Ly6C BV786 (All antibodies were purchased from BioLegend). DAPI was stained separately at 1:5000 dilution for cell viability. Samples were washed, resuspended in FCM buffer (200ul), and were moved to FCM tube before flow cytometry data acquisition in FACS symphony A5 (BDscience). All data were gated and analyzed using FlowJo_v10.10.

### Cytometric bead array (CBA)

To Access cytokine levels, CBA was performed using materials from BD™ CBA Mouse Inflammation Kit (BD Biosciences, 552364), detecting KC, IL-1b, IL-6, IL-10, and TNF-α in accordance to the product guidelines. The staining process was same as CBA, with exceptions to tissue preparation. Tissues were put into Eppendorf tube with 300ul of NP40 lysis buffer (Invitrogen,289375-000) with1:500 dilution of protease and phosphatase inhibitor (GenDEPOT). Metal beads were inserted into a tube containing tissues and tissuelyserII (QIAGEN) was used to lysate the tissues. Chloroform is added (200ul) to the lysate solution, and were centrifuged for 13000rpm for 15 minutes at 4°C. Top forming layer was extracted containing proteins. For Cerebro Spinal Fluid sample, this step was skipped. Then all Assay were performed according to product guidelines. Data was obtained using FACSymphony A5 Flow cytometer. Raw was analyzed using FCAP Array V3 Software and were quantified into graph using Graphpad prism 10.1.

### Neurological Score, Survival curve, & Behavioral test

To access neurological test, Modified Garcia scoring^25^ was performed on Post Operative Day (POD) 1, 3, and 5 with slight modifications on the table (Supplementary Table 1). We excluded lateral turning for the assessment due to possible biased movement surgical suture, as we are performing the test at an earlier stage. At POD 5, any subjects that had a total score below 5 were excluded as a subject for further behavioral test and were only checked for survival rate. Mice were given time for recovery from then up to 4 weeks (POD 27) for recovery. Each mouse was checked every day for survival up to 2 weeks as the highest fatality occurs within this period^26^.

For motor assessment, on POD 28, mice were moved to a behavioral test room 1 hour prior to the rotarod test to get acclimated to the environment. Then, each mouse was placed onto the rotarod and were rotated once. Then, rotarod test was performed to increase rpm at a rate of 4rpm/minute. Latency and final RPM before falling was recorded and analyzed. For cognitive Morris Water Maze (MWM) test, see the supplementary methods section.

### TUNEL staining

To access neuronal cell death, we performed staining within Situ Cell Death Detection Kit, POD (Roche, 12156792910). To observe secondary damage, we selected samples that had blood present in ipsilateral hemisphere of perforation site at POD 3. All harvested mice brain were fixed in 4%PFA in PBS overnight and were made into FFPE. Each section obtained were 6-8 μm. We located targetable area using Allen brain institute 3D Mouse brain atlas (See figure 4A) to obtain section containing motor cortex. To obtain the section of interest, we trimmed from prefrontal cortex to 460 to 480 μm. Once the Lateral ventricles were shaped in accordance with previously viewed image, we obtained sections within 20 μm difference. For obtaining a section with hippocampus, we proceeded additional 400 μm trimming, repeating the procedure.

We performed deparaffination procedure with the following step: Xylene 10 minutes twice, 100% ETOH 10 minute twice, 90% ETOH 10 minute, 50% ETOH 10 minute, Tap water 10 minute. The staining protocol was followed as instructed in the guideline. After staining protocol, samples were counter stained with hematoxylin (Abcam, ab220365). Dehydration step was following deparaffination step in reverse order, with exceptions to Xylene time reduced to 5 minutes. Samples were mounted in Permount mounting medium (Fisher chemical).

All images were slide-scanned using Leica Biosystems-Aperio and were analyzed and obtained using QuPath 0.5.0 Software.

### Immunofluorescence staining

To visualize CCR2, we performed immunofluorescent staining on tissue section or meningeal wholemount mentioned above. First, samples on slides were permeabilized by incubating with 0.3% Triton-X in 1xPBS. Then, samples were washed with PBS twice, 5 minutes each. After washing, samples were blocked using 3% BSA 0.3% Triton-X in PBS for 30 minutes. CCR2 antibody was diluted 1: 200 (abcam, ab273050) in antibody dilution solution (1% BSA 0.1% Triton-X in PBS) and slides were incubated in it over night at 4°C. Next, slides were washed three times with PBS for 5 minutes each, then were incubated in with Alexa 555 anti-rabbit for visualization (Invitrogen, A-21428) diluted at 1:2000 in antibody dilution solution for 1 hour in Room temperature. Then slides were washed with PBS twice and incubated with PBS containing DAPI (1:2000) for 10 minutes. Slides were washed with PBS twice again and were mounted under Vecta-shield antifade mounting medium (Vector lab., H-1900-10). For image acquisition, Leica TCS SP-5 confocal microscope was used for brain section images. For wholemount imaging, we used Benchtop Confocal BC43 (Oxford instrument, Andor) for entire wholemount scanned image. All images were then acquired ImageJ or IMARIS software.

### CCR2 Inhibition

For testing CCR2 antagonist, C57BL/6 mice were used to create SAH model. Approximately 1 hour after the modeling, Mice were given intraperitoneal injection of RS504393 (Medchem, HY-15418) with dose of 9mg/kg, dissolved in 5%DMSO in PBS, for ensured CCR2 inhibition effect based on Tian et al., 2022^27^. After 24 hours, modified Garcia scoring was performed on the mice, and the brain was harvested for FCM data and analysis to verify reduction of infiltrating CD45^hi^ myeloid cells and monocytes.

### Statistical Analysis

All quantified data were statistically analyzed with GraphPad Prism 10.1. All statistical analysis were tested with One-Way ANOVA. The survival curve was statistically analyzed using Mantel-Cox test to see any statistically significant difference.

## Results

### Increase in monocytes occur within the Dura mater of SAH mice during the first 24 hours

First, we wanted to confirm the increase in CCR2^+^ monocytes during the EBI of SAH mice. To mimic rupture and intracranial aneurysm, we established MCA perforation SAH mice model. We confirmed the presence of hemorrhage on the hemisphere ipsilateral to perforation site and observed blood formation near the MCA site (Figure 1A). After establishing the SAH modeling, we wanted to determine if monocyte population increased within meninges after SAH. Meninges have been highlighted as an immune hub for neuroimmune surveillance^28^, and while direct evidence demonstrating increased neutrophils have been present during EBI, evidence showing monocyte and macrophage recruitment has not been well-observed. Using hCD68-GFP reporter mice, which show GFP expression in Macrophage and monocyte lineage, we determined distributions of these myeloid cells within dura mater during early EBI. We generally observed evenly distributed GFP signals throughout the leptomeninges in control mice (Figure 1B). However, after the first 24 hours (POD 1) of SAH, the increase in GFP signal was concentrated within sinuses, regions well known to increase with immune cells during neuroinflammation^29,30^. Especially we saw significant increase in Superior Sagittal Sinus (SSS) (Figure 1C). We further saw an increase in CCR2^+^ signals, along with CCR2^+^ GFP^+^ double positive cells, suggesting an increase in monocyte cells. This evidence confirms macrophages and monocytes are being recruited into the meningeal layers within early EBI of SAH.

**Figure 1.**
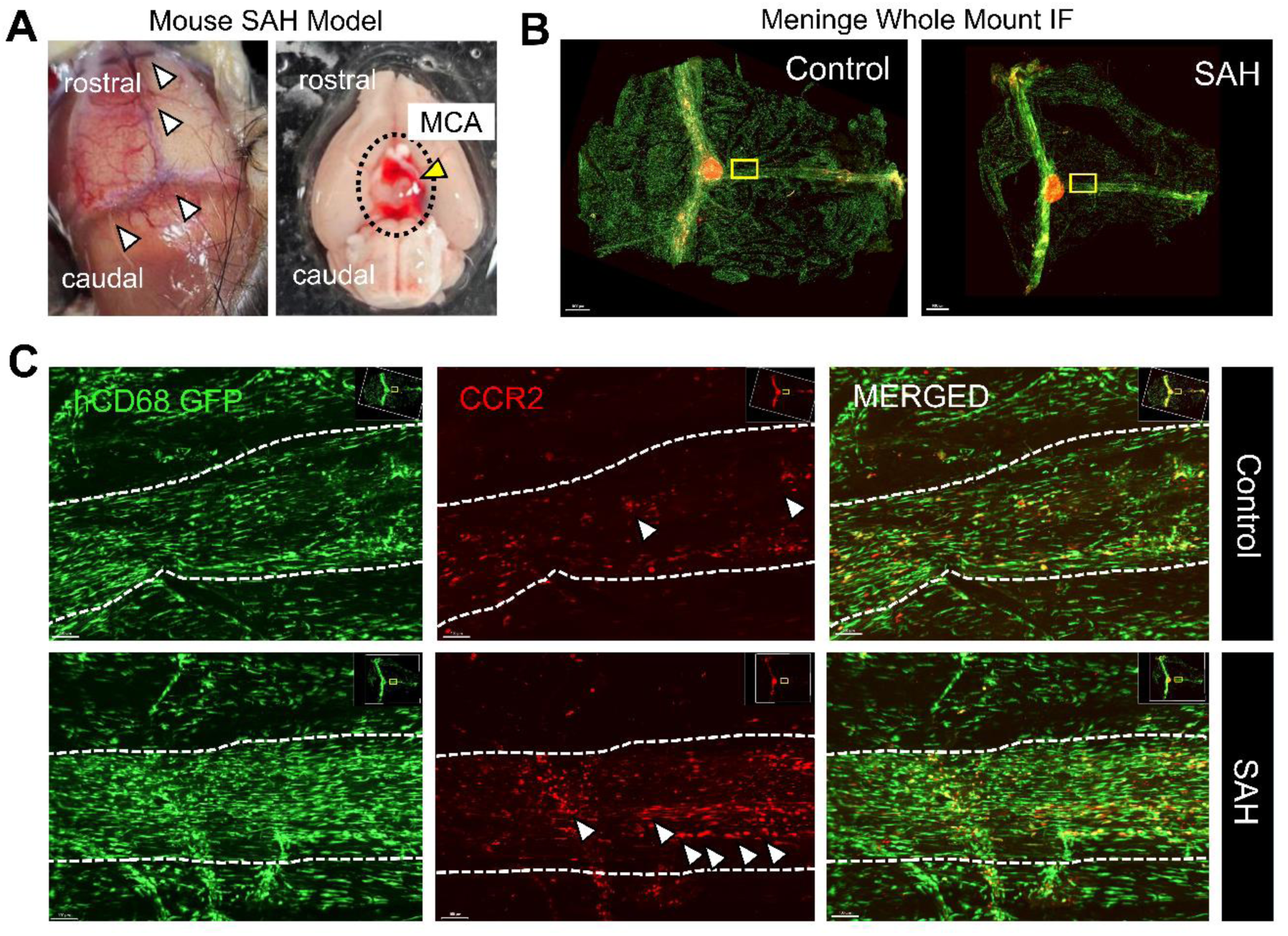
Infiltrating CCR2^+^ cells increase in Meningeal layer during Early EBI of SAH. A) Representative image of SAH mice model confirming hemorrhage in meningeal layer before harvesting brain (left) and bleeding by MCA perforation (right, black dotted circle: Circle of Willis, Yellow arrow: MCA perforation site). B) Meningeal wholemount scanned image of Control and SAH hCD68-GFP mice. Scale bar = 1mm. Yellow square represents zoomed in area of Figure 1C). C) Images of 10x zoom in of SAH and control at superior sagittal sinus, demarcated by the white dotted line. Images of GFP^+^ myeloid cells (Green), CCR2(red), and merged images are shown. White arrows indicate CCR2^+^ cells. EBI: Early Brain Injury, SAH: Subarachnoid Hemorrhage, MCA: Middle Cerebral Artery, CCR2: C, Ctrl: Control *Scale bar =100 μm N=3 per group*.

### Decreased immune infiltration during EBI of SAH mice model in CCR2^-/-^ mice

As we confirmed immune infiltration of Monocyte within meninges during early periods of SAH, we further investigated whether CCR2 inhibition impacted immune cell infiltration. To ensure this, we used CCR2**^-/-^** transgenic mice to create an SAH model and compared immune population with WT SAH model using flow cytometry during the first 72 hours of SAH induction (Figure 2A, B). We first wanted to observe whether the total infiltrating immune cell population (CD45^hi^) had any difference during the early (POD 1) and late (POD 3) period of EBI. Our results showed that during early EBI, compared to wildtype, CCR2**^-/-^**mice had reduced nearly half the amount of infiltrating immune cell population (Figure 2C). However, by late EBI periods, differences in immune population were not present, indicating that immune cell recruitment occurred heavily during the early EBI period of SAH. After observing decreased infiltration, we further analyzed which myeloid cell types are greatly reduced between two SAH groups (Figure 2D). We saw reduced numbers of CD45^hi^ CD11b^+^ Ly6C^hi^ monocyte infiltration on early EBI, which reverted to no difference during late EBI. Both CCR2**^-/-^** and WT SAH models had increased CD11b^+^ Ly6G^+^ neutrophil infiltration during SAH. Also, we did not observe any differences in CD45^hi^ CD11b^+^ Ly6C^-^ macrophage group as well, suggesting that other myeloid cells were only minimally impacted by CCR2 inhibition.

**Figure 2.**
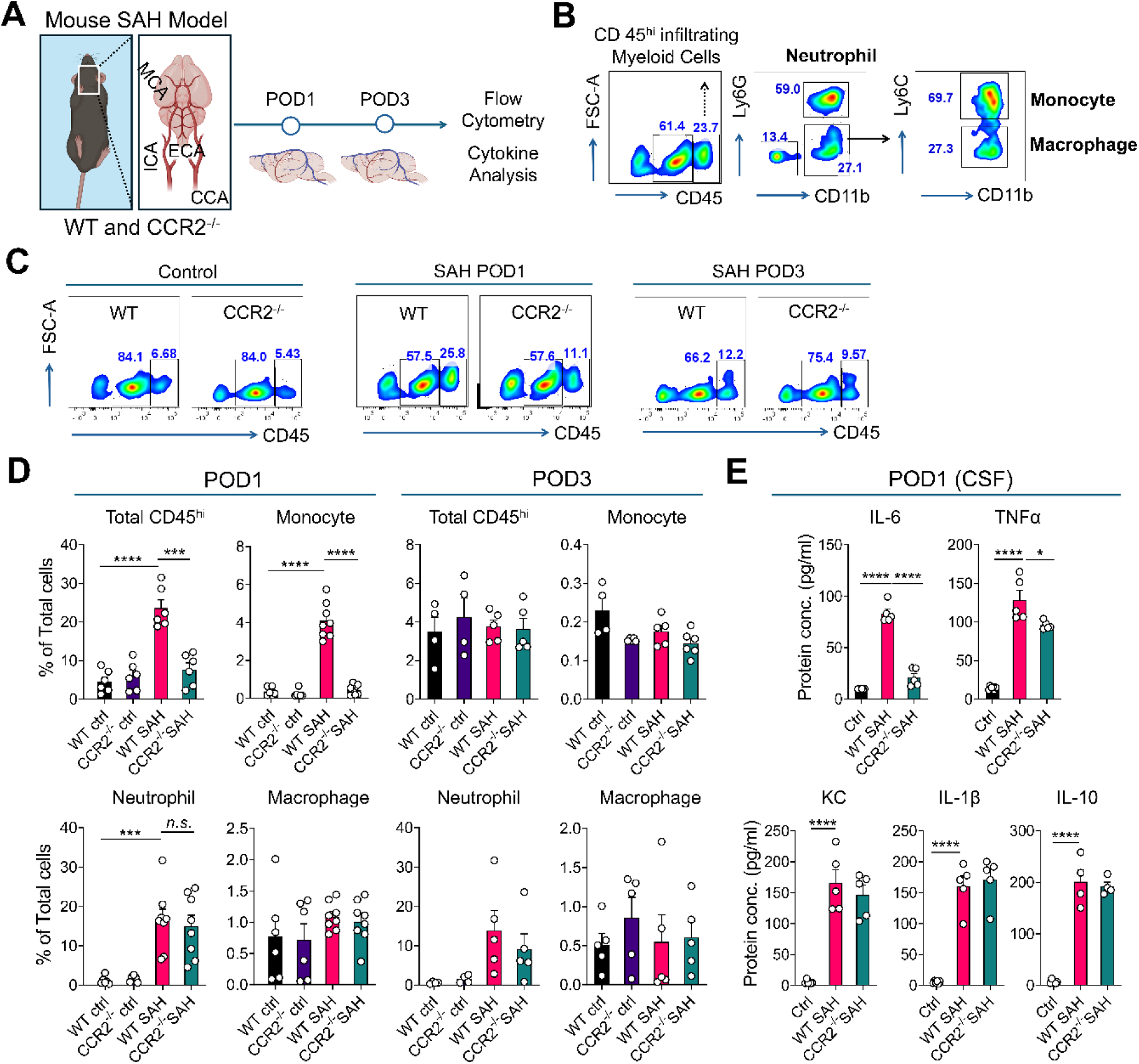
CCR2^-/-^ SAH mice models reduced immune cell infiltration, particularly Ly6C^hi^ monocytes, and reduced CSF IL-6 and TNF-α Cytokines. A) Schematic depiction of SAH modeling to Flow Cytometry performance. B) Gating strategy of Infiltrating myeloid cells, and specific innate immune cells. C) Representative population difference in CD45^hi^ population in Flow cytometry data. D) Quantification of Cells from Flow Cytometry data analyzed at POD1 (Top) and POD3 (bottom). E) CBA data quantified for determining Cytokine concentrations within CSF. POD: Post Operative Day, CSF: Cerebro Spinal Fluid *For POD 1, N=6 per ctrl, N=8 per SAH group. For POD 3, N=4 per ctrl N = 5 per group. One-Way ANOVA, ****= P=<0.0001 *** = P<0.005 ** = P<0.01 * =p<0.05*.

Given the monocyte population difference that occurred during the early EBI, we further investigated cytokine levels to determine differences in neuroinflammation. As we saw no clear difference between CCR2**^-/-^** and WT controls in both cellular and cytokine levels, we compared CCR2**^-/-^** and WT SAH mice with WT controls only. Within the brain lysate, Keratinocyte derived chemokine (KC or CXCL1) and IL-6 had significant difference between CCR2**^-/-^** and WT SAH model (Supplementary Figure 1A). In addition, inflammatory cytokines such as TNF-α or IL-1b and anti-inflammatory cytokine IL-10 had no difference among two SAH models, CCR2**^-/-^** had significantly elevated IL-10 while WT had that of TNF-a compared to its control. Nevertheless, tissue lysate had extremely low level of cytokines; thus, we decided to analyze cytokine level within the Cerebro Spinal Fluid (CSF), as SAH, unlike intracerebral hemorrhage, may not have heme and blood leak into brain parenchyma, which could possibly mean CSF contain accurate concentrations of Cytokine levels (Figure 2E). As expected, the concentrations of all cytokines were elevated close to 10 to 20 times. We continued to see significant differences in IL-6, while detecting some significant reduction of TNF-α cytokine as well. However, CXCL1 concentrations did not show difference within CSF, indicating that CXCL1 signaling, which is important for neutrophil infiltration, may have remained the same. Taking it together, our data suggests that the inhibition of CCR2 signaling results in decreased myeloid cell infiltration within brain and subarachnoid space, mostly monocyte population, and potential reduction of neuroinflammation as shown by decreased cytokine levels such as IL-6 and TNF-α.

### CCR2^-/-^ SAH mice show better recovery and conserved motor score compared to WT

Finding the impact of reduced inflammation during EBI period, we wanted to further determine whether CCR2 inhibition ameliorated the outcome of SAH after survival. Recent research has shown that mice SAH model shows differences in both short term (POD 3 to 7) and long-term recovery (up to POD 28)^31,32^. Based on recent observations, we decided to observe early and late recovery periods of SAH mice model, to see if there’s any difference in its neurological, motor, and memory function (Figure 1 A, Supplementary Figure 2A). First, we determined if there are any survival rate differences for the first 2 weeks (Figure 2B) to see if the inhibition of CCL2 signaling had any impacts in survival of SAH mice models. While the CCR2**^-/-^** did show higher cumulative survival (80%) than WT (53%), there was no statistical significance in terms of survival.

Confirming that CCR2 inhibition has minimal impact in cumulative survival, we further determined to see how neural scoring changed over time. Intriguingly, the modified Garcia scores on POD1, 3, and 5 revealed that CCR2**^-/-^**SAH mice had significantly better scores on all short-term recovery period, showing distinguished improvements in their motor skills on tests, especially on climbing (Figure 3C, Supplementary video 1). We also tested the outcomes of the microglia depleted WT SAH mice model using PLX chow, which showed no recovery compared to the WT SAH (Supplementary Figure 2A), suggesting the presence of microglia is necessary. We further determined long term motor and memory function 4 weeks (POD28) after the SAH (Figure 3D, Supplementary Figure 2B-D). The motor performance was significantly different between WT and CCR2**^-/-^** SAH group, performing as well as the control. Meanwhile, most of the WT SAH groups could not perform as well as either of the group, showing the difference in their prognosis outcome. However, no difference was observed in MWM performance in both learning and probe tests. Our results overall indicate CCR2**^-/-^** SAH mice preserved its motor and neurological function compared to its WT complements, suggesting CCR2 plays a role in recovery after SAH.

**Figure 3.**
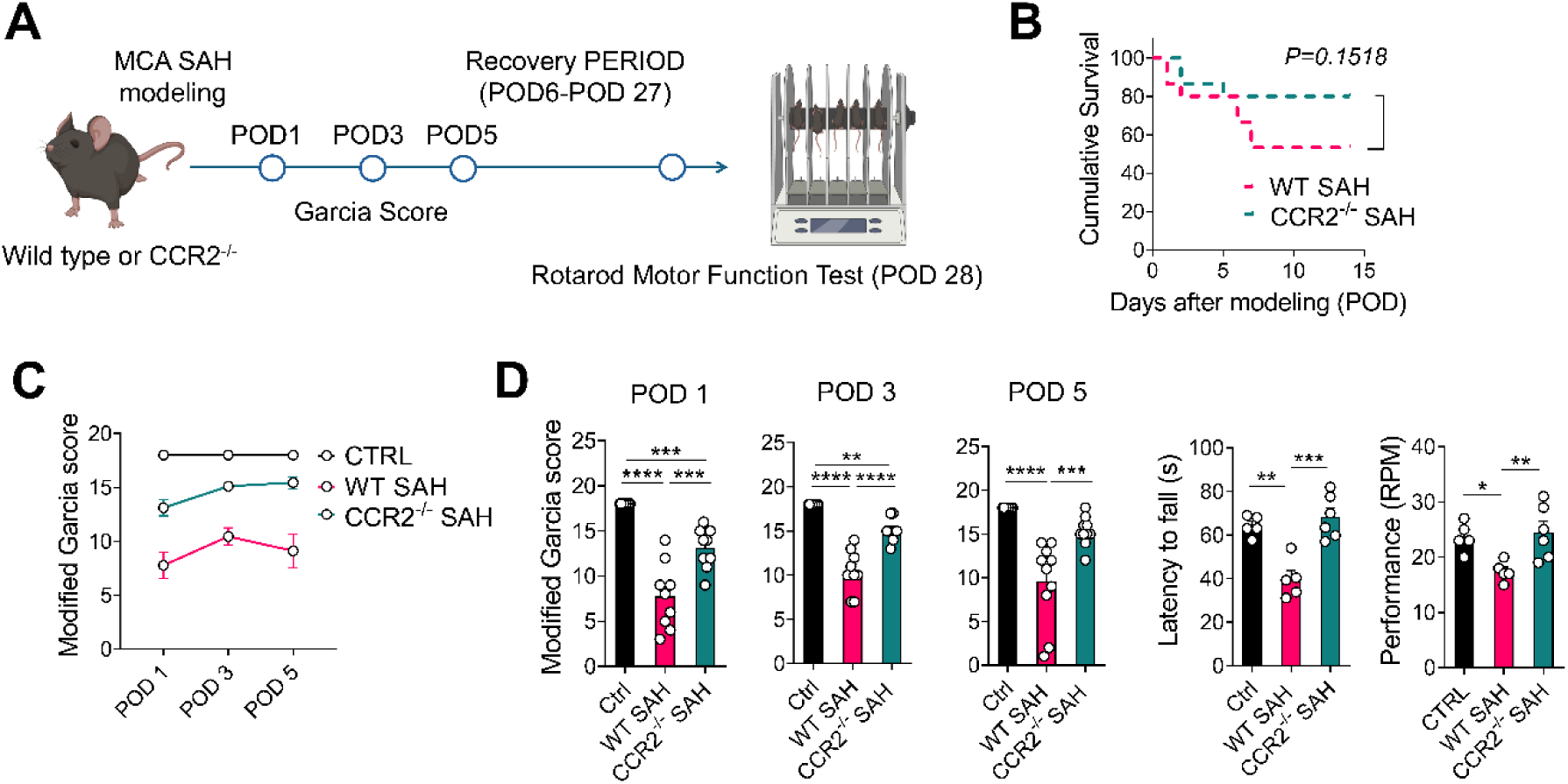
CCR2^-/-^ SAH mice model has better neurological recovery & long-term motor performance. A) schematics of all neurological and behavioral test. B) Survival curve determined up to POD 14. C) Modified Garcia score chart with combined days to show changing average score (left) and individual day (right). D) Data of Rotarod Performance on POD 28 with endurance (Left) and motor performance (Right). *N=8 per group. One-Way ANOVA, ****= P=<0.0001 *** = P<0.005 ** = P<0.01 * =p<0.05*.

### CCR2^-/-^ SAH mice show reduced neuronal cell death compared to WT SAH mice

We have confirmed decreased monocyte infiltration and IL-6 level in CCR2^-/-^ SAH mice and saw its better neurological outcome during recovery. We further wanted to corroborate that inhibiting CCR2 mediated monocyte infiltration resulted in a better outcome. Severe grade of SAH typically results in higher chances of neuronal damage, such as delayed cerebral infarction in SAH patients^33^; given this, we wanted to confirm if CCR2 inhibition resulted in reduced neuronal cell death under severe SAH condition. Based on our flow cytometry data, immune cell infiltration was reducing around POD 3, which during this late EBI period most of cell death would have occurred as it has been known most neuronal death occurs within the first 48 hours of SAH^34^. Specifically, we investigated brain regions relevant to the motor cortex area as the differences were shown in their motor performance. Using Allen Brain Connectivity Atlas (Allen Institute for Brain Science^35^), we obtained cortical sections relevant to this point and performed TUNEL assay to determine cell death (Figure 4A,B). We saw significant differences between the TUNEL^+^ cells among SAH groups in terms of intensity rather than total number of regions. As TUNEL stains for DNA fragmentation, and it could also present different cellular apoptotic stage, or amount of DNA damage given to these cells^36^. When observing the brain within motor cortex area of the brain specifically, we noticed differences within intensity of TUNEL (Figure 4C); while WT SAH had strong TUNEL (TUNEL^hi^) signals, CCR2^-/-^ SAH mostly had weaker TUNEL (TUNEL^lo^) or no TUNEL. We also stained for IBA1^+^ microglia within the cortex to determine if there were any differences in its activation morphology, however no difference was found (Supplementary Figure 3A). Intriguingly, when we also analyzed dentate gyrus (DG) to see differences with TUNEL signals, no visible differences were present, with exception of faint signals from SAH WT, indicating that DG has a minimal impact within this model in terms of neuronal cell death during the EBI period. When we quantified total TUNEL^+^ cells within the brain, we found that CCR2**^-/-^** SAH mice group had approximately 75% reduced TUNEL signals compared to WT SAH mice group (Figure 4D) on average. Taken together, our results suggest that CCR2**^-/-^** improved neuronal cell death outcome under severe SAH condition in comparison WT mice during late EBI period.

**Figure 4.**
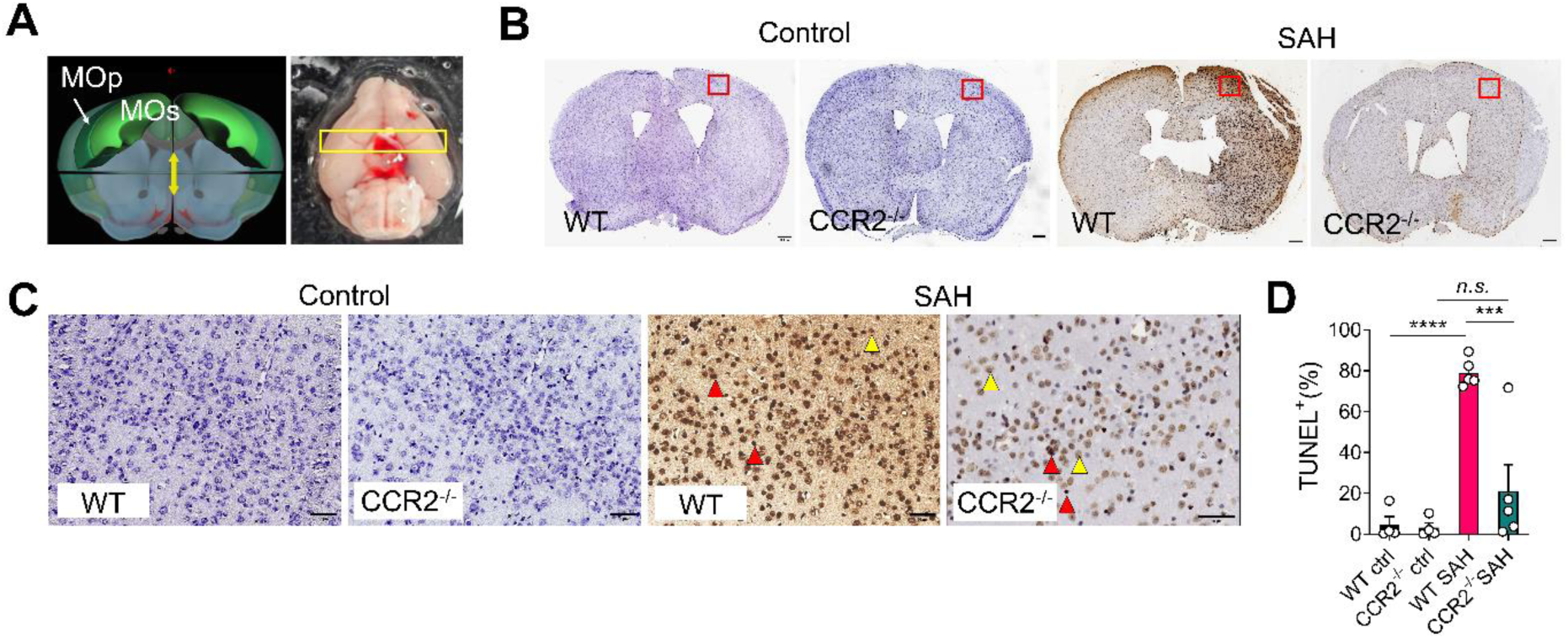
TUNEL Assay shows CCR2^-/-^ SAH mice model has reduced neuronal cell death. A) Selected area for TUNEL staining. We used *ALLEN 3D BRAIN ATLAS* **(left)** for accurate plane of section where motor cortex was present, indicated by the yellow box drawn on actual sample (right). B) representative image of TUNEL staining for each group. Red squares indicate the zoomed in region of motor cortex shown in figure C. Scale bar = 500 μm C) zoomed in motor cortex area of the slide scan in Figure B. Red arrows indicate examples of TUNEL^hi^ cells, while yellow arrows indicate examples of TUNEL^low^ cells. Scale bar = 50 μm. D) quantified TUNEL staining with total count proportion (left) and classified staining (right). *One-Way ANOVA,****= P<0.0001 *** = P<0.005 ** = P<0.01 * =p<0.05 N=3 for each Ctrl, N=5 for each SAH group*.

### CCR2 inhibition before EBI period in WT SAH mice model ameliorates recovery burden

As we have confirmed that CCR2**^-/-^** SAH mice model showed improved neurological and motor behavior after SAH. We wanted to see drug induced CCR2 signaling inhibition before EBI period could ameliorate neurological deficits after SAH (Figure 5A). We compared Garcia scoring and immune cell composition at POD 1, as our data confirmed infiltrating immune cells within this period to be the highest. Garcia score showed WT SAH model treated with CCR2 inhibitor displayed higher neurological scoring compared to vehicle and intriguingly showed similar improvements in behavioral performance as previously observed in CCR2**^-/-^** SAH model (Figure 5B, Supplementary Video 2). We also confirmed that these mice had reduced myeloid cell infiltration as well as decreased monocyte, suggesting drug treatment worked successfully. We wanted to further validate the induction of CCR2 decreased; when observing parenchyma near the MCA perforation site, we observed decreased CCR2 signaling compared to Vehicle group, further confirming that the drug treatment successfully inhibited CCR2^+^ cell infiltration near the perforation area (Figure 5D). Taking all results together, CCR2 inhibition resulted in amelioration of SAH progression especially during EBI period, which may have impact in disease progression, conserving motor damage.

**Figure 5.**
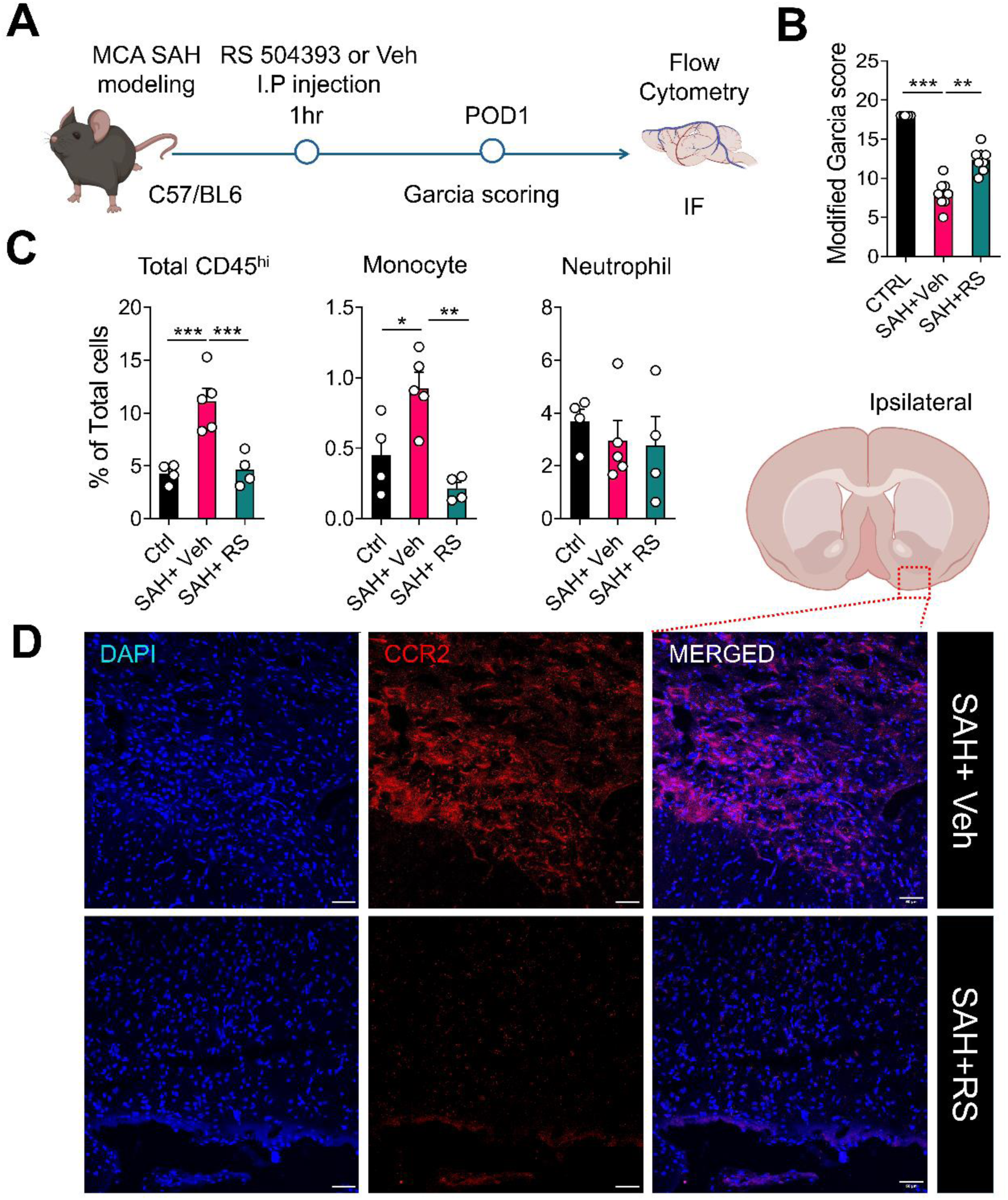
CCR2 antagonist treatment at the beginning of EBI ameliorates WT SAH mice model. A) Schematics of CCR2 antagonist administration experiment after SAH induction. B) Graph representing Garcia scoring of SAH mice model on POD1. C) FCM analysis data of infiltrating innate immune cell population on POD1. D) representative Immunofluorescence imaging showing CCR2+ cells between SAH mice treated with either Vehicle or CCR2 antagonists. Area of region depicted by Orange square shown in left figure. Red X indicates ipsilateral side of perforation site. *Figure B and C, N= 4 to 5 per group. For figure D, N= 4 each group. Scale bar = 50 μm. One-Way ANOVA, ****= P=<0.0001 *** = P<0.005 ** = P<0.01 * =p<0.05*.

## Discussions

SAH holds its unique position as its pathogenesis begins outside brain parenchyma, and integration of well-known immunological principles into such specific condition is required^37^. While roles of innate immune cells have been well established in different cases of autoimmune or neurodegenerative murine models^38,39^, in the events of stroke, roles of infiltrating and resident immune cells remain controversial, and requires further investigations^40^. To investigate the roles of infiltrating innate immune cells, our present study used CCR2**^-/-^** transgenic mice model to generate SAH model that inhibit infiltrating monocytes and compared it with WT SAH model. We showed decreased overall infiltrating CD45^hi^ populations, with main difference in the Ly6C^hi^ monocyte population, suggesting monocytes to be the main innate immune population recruited during EBI phase of SAH. We also observed reduced inflammatory of certain cytokine levels such as IL-6 and TNF-α in CSF, which suggests possibly attenuated neuroinflammation. In addition, the behavioral outcome had significant improvements, in both Garcia score and motor test, although no significant differences were observed in cognitive deficits. Furthermore, using CCR2 inhibitor, we also found improvements in WT SAH model during the early EBI in its neurological outcome. Taking everything together, our study shows strong evidence that CCR2 inhibition may reduce pro-inflammatory roles of infiltrating monocytes, which supports CCR2 as a therapeutic target for improved neurological outcomes for surviving SAH patients.

The observation of decreased myeloid population seemed evident, as many studies suggest CCR2 signaling as a key mechanism in recruitment of monocytes. The intriguing point is that we also observed reduced cytokine levels of IL-6 and TNF-α, while no differences were found in levels of IL-1b. This could be those types of cytokines induced may be dependent on innate immune cell types. For instance, some studies have suggested microglia as the critical source of IL-1b production during early phases of SAH^41,42^. Since CCR2 does not greatly impact resident macrophages like microglia within SAH context, microglial inflammatory response may not have changed, which can explain our result. However, the decreased monocyte population, which occurred along with reduced TNF-α and IL-6 level, may be critical to the sustained inflammatory damage during EBI phase. SAH results in an increase of IL-6 level in CSF due to neuroinflammation, which is considered as one of the biomarkers of extensive prognosis of SAH, yet the complex roles of IL-6 are not fully understood^37,43,44^. Perhaps it is possible that CCR2^+^ infiltration may be a key population in producing these cytokines, and reduced cell recruitment potentially ameliorated neuroinflammation. Our study could not address whether CCR2 inhibition had impacts on other neuroprotective effects, or whether it influenced Blood brain barrier permeability. Further investigations such as measurement of NF-κB and microglial activation under CCR2 inhibition is necessary to clarify and fully characterize the anti-inflammatory effect. It would also be interesting to see whether decreasing IL-6 and TNF-α using antagonists could produce similar neurological outcomes.

Differences in neuronal death using TUNEL assay displayed considerable differences between two SAH groups. As stated at the beginning, physical pathological stress like increased ICP cannot be controlled by CCR2 inhibition, which would indicate why CCR2^-/-^ SAH model still maintained neuronal death. However, possibly reduced neuroinflammation may have had neuroprotective effects from other possible risks during EBI. Recent studies showed early anti-inflammatory drug like ibuprofen resulted in potential benefits in overall outcomes, including attenuation of neuronal damage and decreased vasospasm ^45–47^, although therapeutic effects were limited. It is likely that the EBI time window is critical, and unspecific immune suppression may be of issue. In an immunological perspective, repairment of blood vessel ruptures requires pro-inflammation phase^48,49^; uncontrolled immune suppression through may result in adverse effect, and it is likely that balance between pro- and anti-inflammatory is the key to controlling any additional damage. In this sense, rather than aiming for complete immune suppression, selective cytokine suppressions like anti-TNF-α or anti-IL-6 may be beneficial. In our study, the inhibition of monocyte recruitment led to selectively reduced cytokine level, indicating that these cells are potentially key immune cells in causing increased pro-inflammatory reaction, breaking the balanced recovery phase. Further study investigating whether monocytes are the key origins of specific types of cytokines would be important to confirm discrete target which may be of good therapeutic use for SAH prognosis outcome.

Our SAH perforation model showed a remarkably positive neurological outcome during recovery in both CCR2**^-/-^** SAH mice model and in drug administration of CCR2 inhibitor. The results shown could be due to the reduced cytokine level mentioned above, which ultimately led to a milder inflammatory response. One of the most intriguing parts about our neurological outcome is that CCR2**^-/-^** SAH mice model had better performance and recovery during EBI period, indicating its beneficial effects in the short term as well. Our results also demonstrated that the CCR2 inhibition in WT SAH model had improvements. Recent clinical study has confirmed monocyte-related ratio is significant predictors of patient outcomes of SAH^50^. In addition to this, our study serves to not only corroborate the importance of monocyte infiltration towards subarachnoid space, but early intervention of CCR2 is beneficial in SAH outcome. One concern is that MCA perforation model is limited for controlling the severity of SAH, we were unable to investigate whether this is applied to all SAH mice model with different grades. Further investigation in severity dependent monocyte inhibition is universally beneficial to patient outcome may be necessary.

In conclusion, we believe that CCR2 inhibition during EBI reduces monocyte infiltration during SAH, resulting in reduced cytokine levels, neuronal cell death in motor cortex, and improved neurological outcomes. We believe our study supports the potential of targeting CCR2 during EBI as an intervening therapeutic strategy, which could benefit surviving patient’s outcome.

---

**Supplemental Figure 1.**
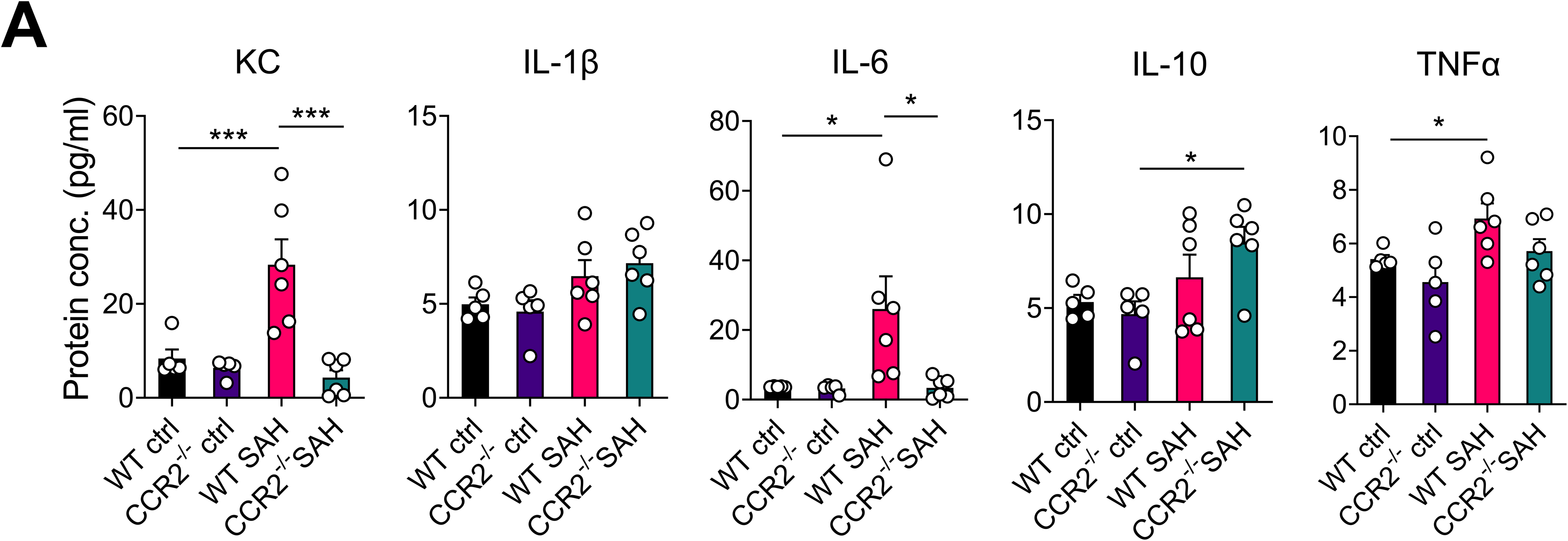

**Supplemental Figure 2.**
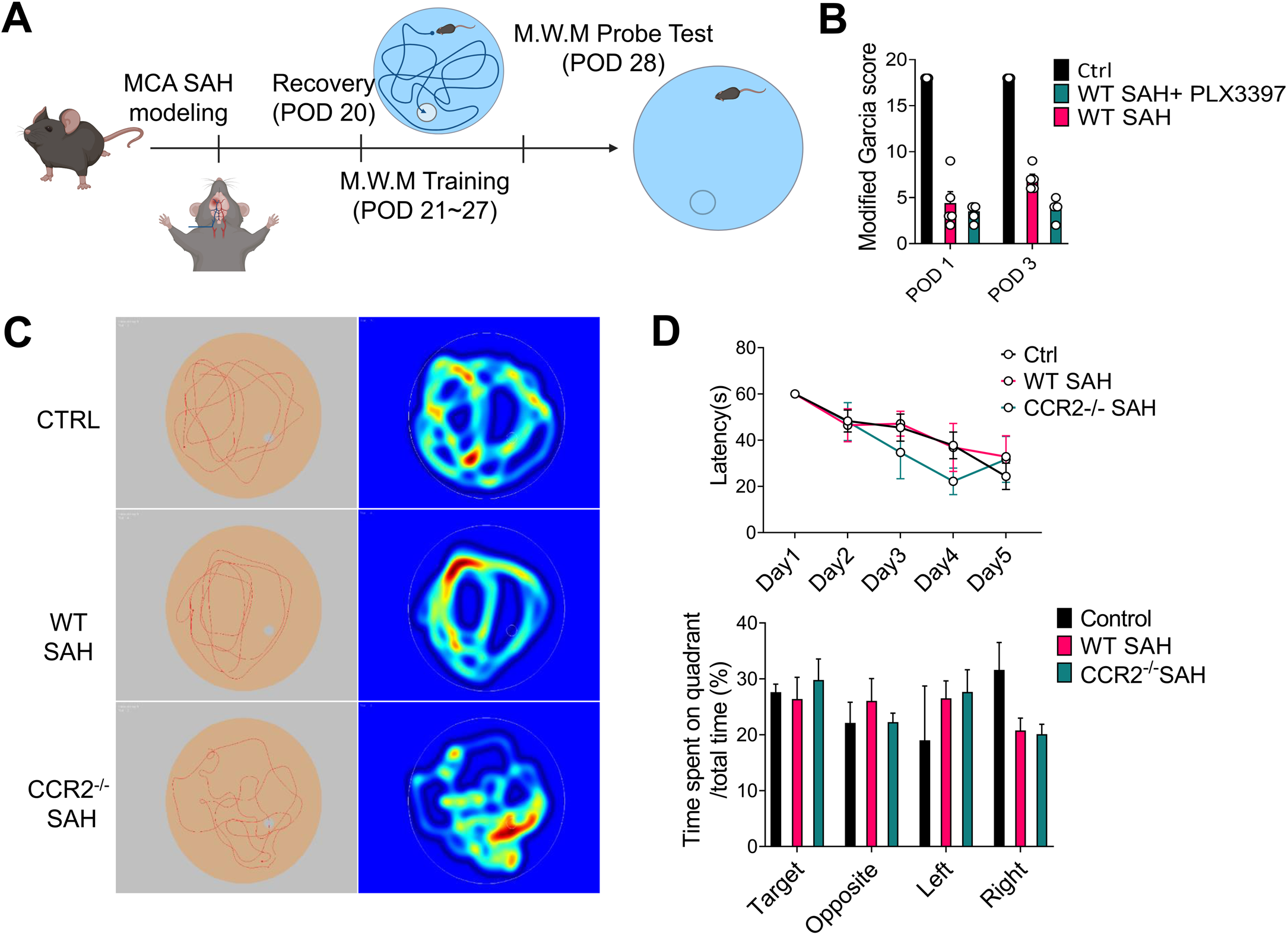

**Supplemental Figure 3.**
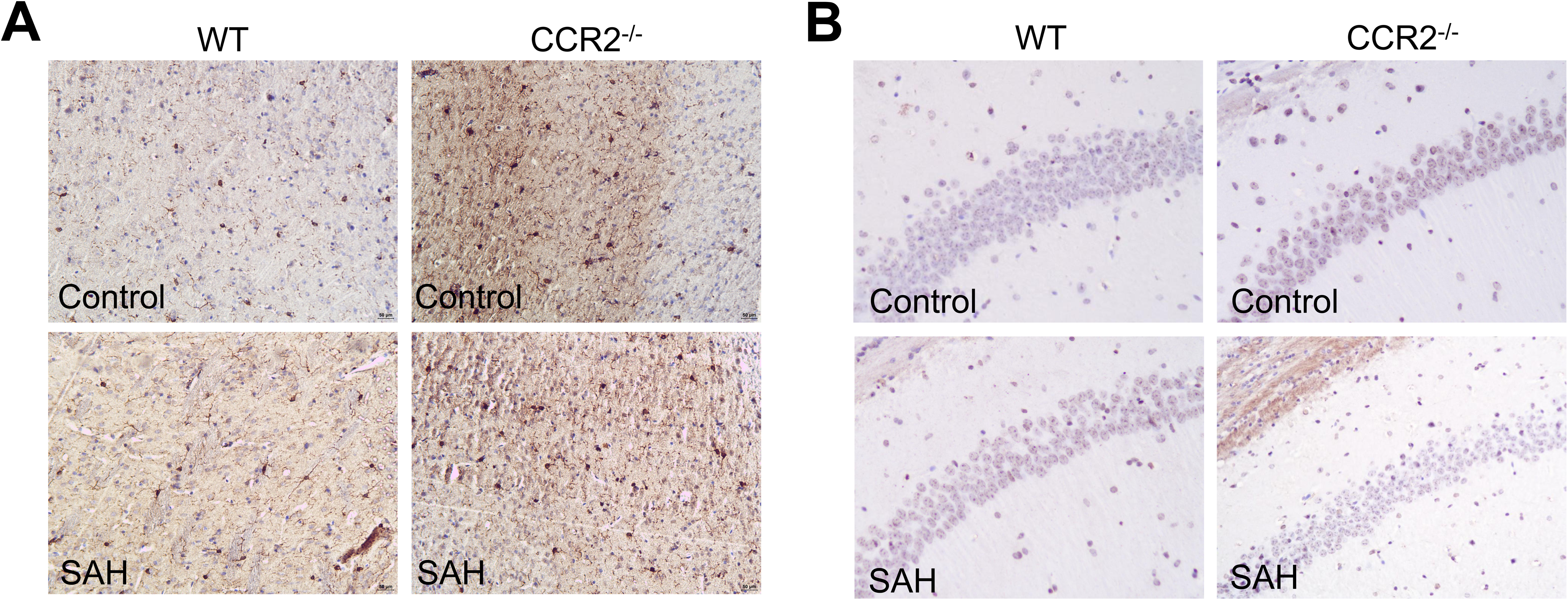

## Abbreviations

EBI: Early Brain Injury (Period)
SAH: Subarachnoid Hemorrhage
DCI: Delayed Cerebral Ischemia
CCL2/MCP-1: C-C Chemokine Ligand 2/ Monocyte Chemoattractant Protein-1
CCR2: C-C Chemokine Ligand Receptor 2
KC/CXCL1: Keratinocyte-derived Chemokine/ Chemokine (C-X-C motif) ligand 1
TNF-α: Tumor Necrosis Factor-alpha
IL-1β: Interleukin-1 beta
IL-6,10: Interleukin 6,10
MCA: Middle Cerebral Artery
POD: Post-Operative Day

